# Integration of Molecular Interactome and Targeted Interaction Analysis to Identify a COPD Disease Network Module

**DOI:** 10.1101/408229

**Authors:** Amitabh Sharma, Maksim Kitsak, Michael H. Cho, Asher Ameli, Xiaobo Zhou, Zhiqiang Jiang, James D. Crapo, Terri H. Beaty, Joerg Menche, Per S. Bakke, Marc Santolini, Edwin K. Silverman

## Abstract

The polygenic nature of complex diseases offers potential opportunities to utilize network-based approaches that leverage the comprehensive set of protein-protein interactions (the human interactome) to identify new genes of interest and relevant biological pathways. However, the incompleteness of the current human interactome prevents it from reaching its full potential to extract network-based knowledge from gene discovery efforts, such as genome-wide association studies, for complex diseases like chronic obstructive pulmonary disease (COPD). Here, we provide a framework that integrates the existing human interactome information with new experimental protein-protein interaction data for FAM13A, one of the most highly associated genetic loci to COPD, to find a more comprehensive disease network module. We identified an initial disease network neighborhood by applying a random-walk method. Next, we developed a network-based closeness approach (C_AB_) that revealed 9 out of 96 *FAM13A* interacting partners identified by affinity purification assays were significantly close to the initial network neighborhood. Moreover, compared to a similar method (local radiality), the C_AB_ approach predicts low-degree genes as potential candidates. The candidates identified by the network-based closeness approach were combined with the initial network neighborhood to build a comprehensive disease network module (163 genes) that was enriched with genes differentially expressed between controls and COPD subjects in alveolar macrophages, lung tissue, sputum, blood, and bronchial brushing datasets. Overall, we demonstrate an approach to find disease-related network components using new laboratory data to overcome incompleteness of the current interactome.

## Introduction

Chronic obstructive pulmonary disease (COPD) is the third leading cause of death worldwide, and recently it was estimated that COPD cases in developed countries would increase by more than 150% from 2010 to 2030 ^1-4^. Furthermore, similar to other complex diseases, it has been challenging to identify systematically the likely multiple genetic risk factors for COPD. Genome-wide association studies (GWAS) can identify specific genetic loci consistently associated with disease in an unbiased manner and have reported hundreds of associations between complex diseases and traits ^5-7^. However, for the vast majority of such genome-wide “hits”, specific causal mechanisms remain uncertain. There is increasing evidence supporting the hypothesis that the onset and progression of complex diseases like COPD arise from the interplay between a number of interconnected causative genes in a manner compounding the effects of any one variant^8-10^. Indeed, integrating GWAS data with molecular interaction networks and gene expression information facilitates a better understanding of disease pathogenetic mechanisms^10-14^. A variety of approaches have been developed to infer relationships between genes showing genome-wide significant evidence of association within the human interactome—the comprehensive set of molecular relationships between cellular proteins ^14-17^. For example, we showed that a disease network module is enriched for disease susceptibility variants in asthma^10^. A GWAS of inflammatory bowel disease used DAPPLE, which is based on the observation that truly causal genes tend to link to each other in the human interactome, to prioritize potential disease candidates^18^. Since combinations of genetic alterations associated with a disease might affect a common component of the cellular system, module-centric approaches might be helpful in finding the disease-related components in the interactome ^13,19^. Yet, the output of these approaches can be strongly influenced by (i) the incompleteness of the pre-specified interactome (false-negative results), and (ii) false-positive errors in the interactome. The impact of the incompleteness could result in failure to identify network relationships for genes implicated by GWAS. Thus, integrating the module-centric approach with targeted interaction analysis (e.g., pull-down assays) of GWAS genes might be helpful in discovering the functional relationships of these genes with a disease of interest. In this work we combine new experimental protein-protein interaction data with the existing human interactome to enhance our understanding of the genes involved in COPD. The objective relies on the “local impact hypothesis,” which assumes that if a few disease components are identified, other components are likely to be found in their vicinity of the human interactome ^10,12^. Moreover, if a disease gene is not mapped in the interactome, it is possible that its neighbors detected by targeted interaction analysis might indicate it’s biological function. Hence, we first identify the disease-related network neighborhood including known COPD disease genes (seed genes) in the interactome by applying a degree-adjusted random-walk algorithm^20^ (DADA), which is a guilt-by-association approach. Next, we test whether experimentally determined links (pull-down assay) for a single, consistently associated COPD gene (FAM13A) not mapped on the human interactome could enhance our knowledge about functional implications of FAM13A in COPD pathogenesis. The approach first aggregates the network neighborhood around the COPD ‘seed’ disease genes using DADA ^20^. Further, to define a boundary of the disease network neighborhood, we use the sub-genome-wide significant association signals from the COPD GWAS (**Figure 1**). This step helps to find enrichment of moderate p–value signals associated with those neighboring genes that are in the proximity of the seed genes. We hypothesized that combining experimental interaction data with the existing human interactome would develop a more comprehensive disease network module for COPD. To test this hypothesis, we derive a novel network-based closeness approach (C_AB_) to predict *FAM13A* partners significantly close to the initial COPD localized neighborhood. Overall, our approach enhances our understanding about the COPD disease network module and predicts new candidate genes and pathways influencing COPD pathogenesis.

**Figure 1:**
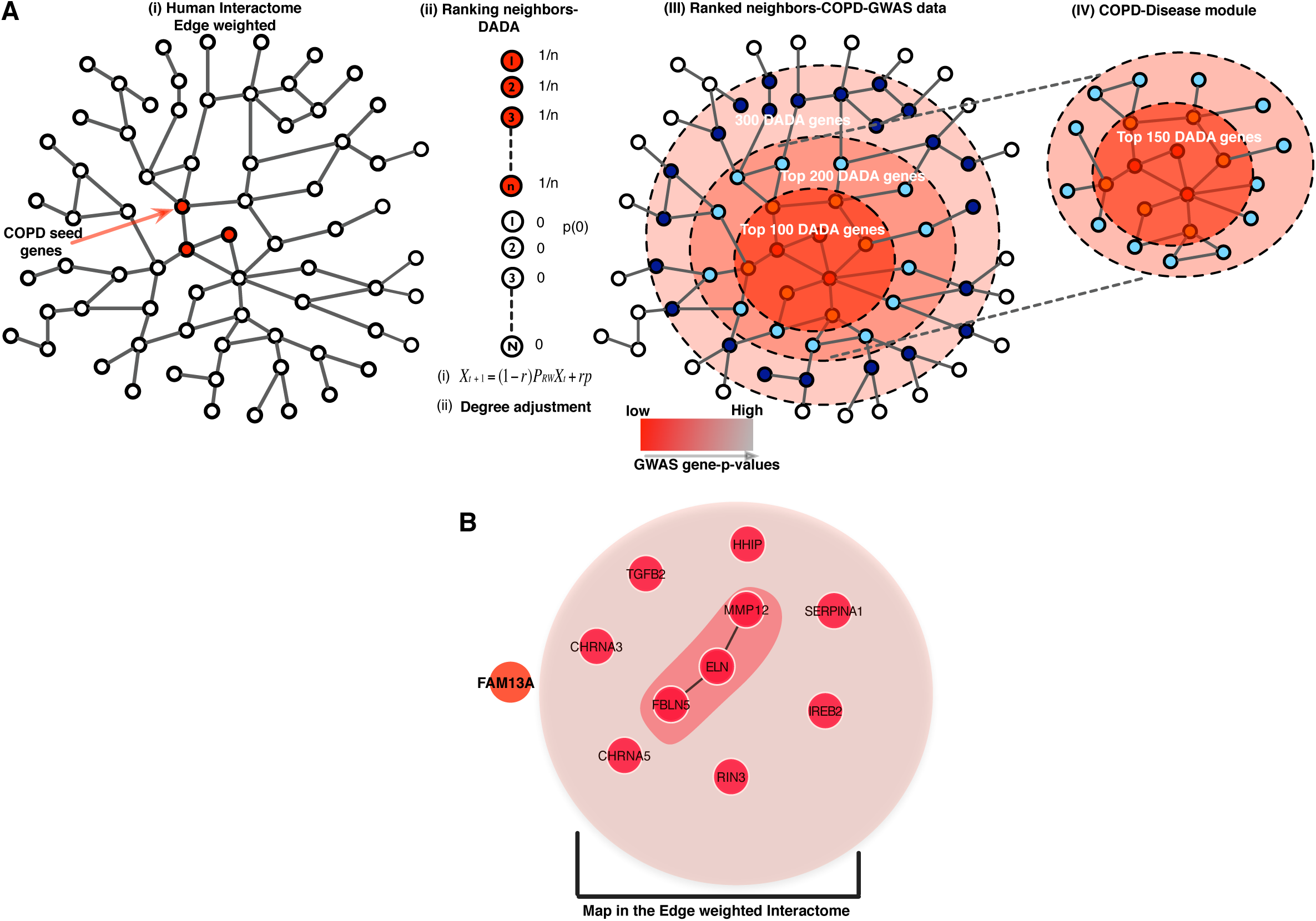
Overview of the approach to identify the COPD disease network module by using the edge-weighted interaction network. First, we applied the Degree-Aware Disease Gene Prioritization (DADA) algorithm and we prune the DADA results by integrating COPD GWAS data. ***A.*** Workflow describing the method. **B**. Among the 11 high confidence COPD seed genes, 10 were mapped on the human interactome, with 3 of them being directly connected.

## Results

### Building an initial COPD network neighborhood using the Degree-Aware Disease Gene Prioritization approach (DADA)

The disease module hypothesis postulates that disease susceptibility genes should form one or a few large connected components in a well-defined neighborhood of the human interactome^10,12^. Selection of the seed genes strongly influences the interpretation of such a module-centric approach, and therefore we restricted our analysis to only high-confidence COPD disease genes from GWAS and Mendelian syndromes (Figure 1B). To avoid bias toward including highly connected genes in the network neighborhood, we implemented the random walk-based DADA approach, which provides statistical adjustment models to remove the bias with respect to degree of the genes^20^. Since DADA provides ranking to all of the genes in the human interactome, we defined the boundary of the disease network neighborhood by integrating additional genetic signals from COPD GWAS (not reaching traditional p-value thresholds for genome-wide significance) (**Supplementary figure 1)**. We first generated a single genetic association *p*-value for each gene in the interactome using VEGAS with the default all snps test ^21^, and then plotted p-values of the added DADA genes vs. the background p-value distribution (**Figure 2A**). After the addition of 150 genes, the genetic association p-value of added genes reached a plateau (**Figure 2A**) and the connected components among the 150 genes were defined as the ‘initial network neighborhood’. At this threshold, we found eight seed genes in the largest connected component (LCC) of size 129 genes, and the other two seed genes were part of two small components of sizes 17 and 4, respectively (**Figure 2C**). Indeed, the LCC of 129 genes was found to be significant compared to the largest connected component that would emerge by chance if the 129 genes were placed randomly (10,000 times) in the human interactome (Z-score =27, p=<0.00001, **Figure 2B**). Overall, these three components constitute the COPD localized neighborhood with 140 DADA genes plus 10 original high-confidence COPD seed genes. We compared our results with the Disease Module Detection (DIAMOnD) algorithm, which identifies the disease neighborhood around a set of known disease proteins based on the connectivity significance ^22^. Interestingly, we found a significant overlap between DADA and DIAMoND output (**Supplementary figure 2**), indicating that the results are consistent using a different network-based approach.

**Figure 2:**
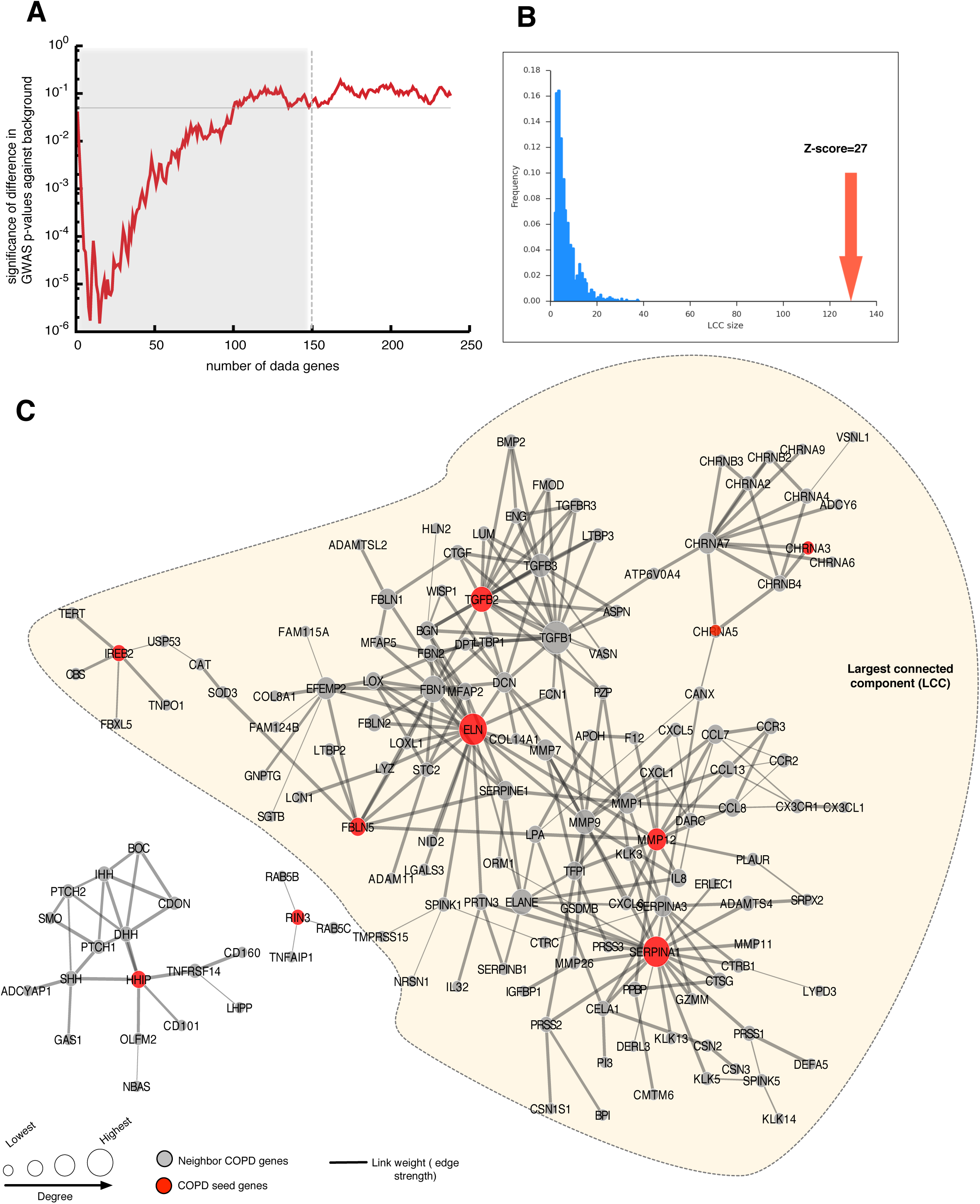
Initial COPD disease network neighborhood. **A**. GWAS p-values of the added DADA genes vs. the background p-value distribution (150 gene cut-off). **B**. Z-score significance of the largest connected component (LCC). **C**. COPD localized network neighborhood of 140 DADA genes and 10 seed genes distributed in three components.

The 10 COPD seed genes that were part of the initial network neighborhood included: *IREB2, SERPINA1, MMP12, HHIP, RIN3, ELN, FBLN5, CHRNA3, CHRNA5*, and *TGFB2* (**Figure 2C**). Since one of the key genes identified by COPD GWAS, *FAM13A*, was not mapped in the human interactome, we tested whether specific interacting partners of *FAM13A* could reveal new knowledge regarding this particular gene in COPD.

### FAM13A pull down assay^23^

*FAM13A* contains a Rho GTPase-activating protein-binding domain; it inhibits signal transduction and responds to hypoxia. Recent work by our research group indicates that *FAM13A* is involved in WNT/beta catenin pathway signaling^23^. *FAM13A* was not mapped in the edge-weighted human interactome (ConsensuspathDB) and moreover, no edges were reported in Rolland et al (2014)^24^ high-quality human binary protein-protein interactions and BioGRID interaction data (2014)^25^. Thus, we performed a pull-down assay using affinity purification-mass spectrometry, which identified 96 interacting partners of FAM13A. We measured the likelihood of having a protein with at least 96 interacting proteins in the interactome. Among 14,280 genes in the interactome, 581 genes had a degree of 96 or greater (*P*(*k* ≥ 96) = 0.04), suggesting that FAM13A is a relatively highly connected protein in the interactome (**Supplementary figure 3A)**. Further, we tested whether the FAM13A interacting partners are closer to each other within the interactome than a same-sized set of randomly selected proteins. Based on 10,000 simulations, we observed significant closeness (Zscore= −9.685) among FAM13A partners (**Supplementary figure 3B**). This indicates that even if FAM13A partners are not directly interacting, they might be involved in a similar biological process because of their close proximity to each other. We found that none of the 96 FAM13A interacting partners were among the COPD localized neighborhood that we had created with DADA.

**Figure 3:**
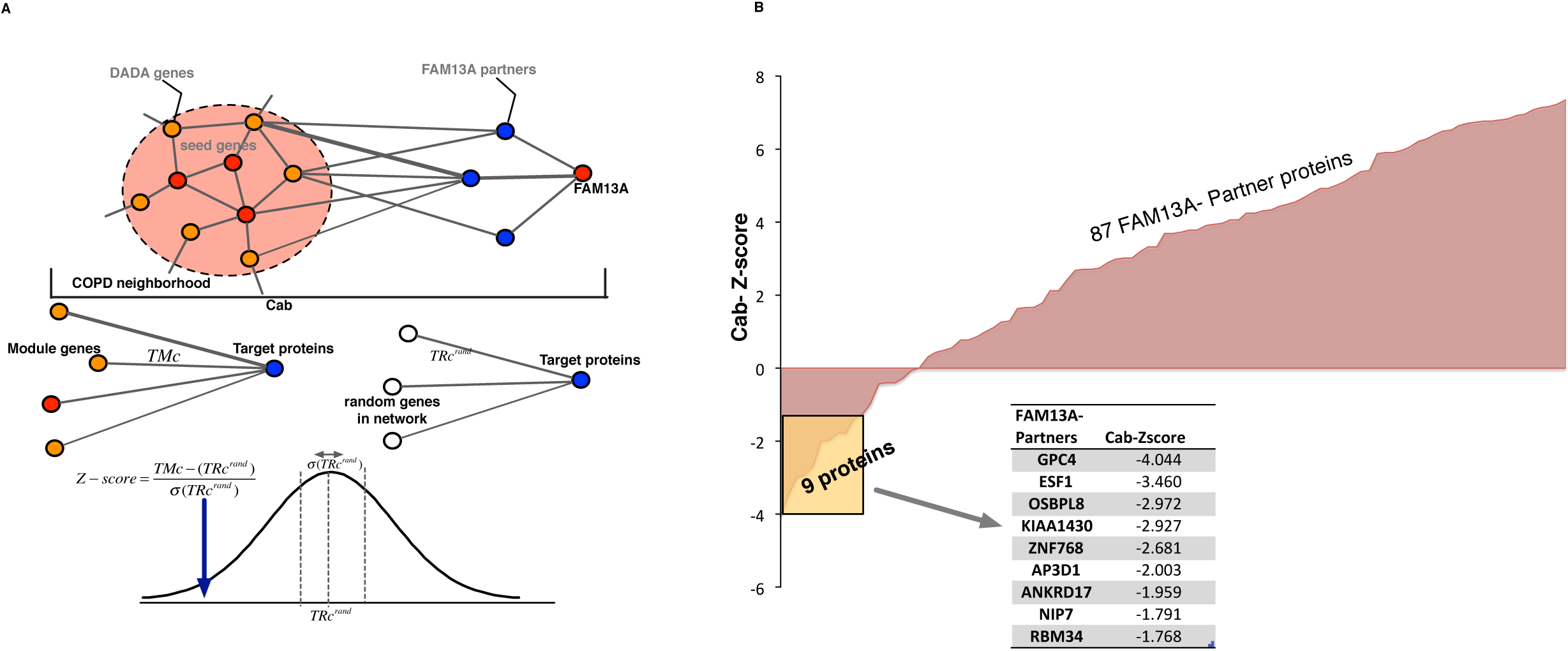
Network-based closeness of *FAM13A* partners to COPD disease network neighborhood. **A**: Illustration of the network-based closeness measure (⟨C_AB_⟩) for FAM13A partners to COPD disease network neighborhood. We calculate the mean shortest distances between ⟨CA⟩and ⟨CB⟩and compare it with the random selection of same number of nodes. **B**: The closeness significance of 96 FAM13A partners to COPD disease network neighborhood.

### Topological distance between the COPD neighborhood proteins and FAM13A interacting proteins in the interactome

Given the substantial incompleteness of the current human interactome ^12^, it is difficult to conclusively determine whether the COPD disease network neighborhood would directly connect to interacting partners of FAM13A, as a single missing link might have disconnected FAM13A from the COPD localized neighborhood. Hence, we computed a network-based closeness metric (C_AB_) that compares the weighted distance between FAM13A partners (*A)* and proteins in the COPD localized network neighborhood (*B)* to random expectation in order to compute the Z-score (see methods and **Figure 3A**). With a Z-score significance threshold of −1.6 (p<0.05), we found 9 genes significantly close to the COPD localized neighborhood in the human interactome and 87 genes that were not significant (**Figure 3B**). The 9 genes with significant closeness to the COPD localized neighborhood were: GPC4 (Z=-4.04), ESF1 (Z=-3.46), OSBPL8 (Z=-2.97), KIAA1430 (−2.93), ZNF768 (Z=−2.68), AP3D1 (Z=-2.00), ANKRD17 (Z=-1.96), NIP7 (-Z=1.79) and RBM34 (Z=-1.77).

### Comparison with the Local Radiality (LR) method

We compared the C_AB_ results with the Local Radiality (LR) method that utilizes topological information (i.e., shortest path distance) to predict the proximity of dysregulated genes to corresponding drug targets ^26^. In our case, we measured the closeness of FAM13A partners (96 genes) with the COPD disease neighborhood (150 genes) by applying the LR method. In C_AB_ the confidence scores of the edges play an important role to either shorten or increase the distances. Thus, to carefully claim that a gene is close to the COPD network neighborhood, we not only ensured that the gene is topologically close to the neighborhood but also considered the strength of each interaction based on different sources of evidence for the existence of such a path. As compared to top C_AB_ genes, the nine highest score genes by LR were enriched in hubs. As a consequence, the average degrees <k> between these two methods were significantly different (P=0.0004, Mann– Whitney U test) (**Supplementary figure 4**). The hubness criterion helped us discriminate between the results from these two approaches. This seems pragmatic, as the low degree genes might be more likely to be involved in a local biological process than those high degree genes representing global molecular pathways. Furthermore, it has been proposed that highly connected superhubs perform the most basic biological functions (evolutionarily early), with the more specialized functions (evolutionarily late) being performed by the peripheral genes. Thus, C_AB_ helps to predict the FAM13A partners that might be involved in more specialized biological functions (low degree genes) related to COPD pathogenesis. Furthermore, it has also been observed that changes in gene expression predominantly occur in the genes (nodes) with low connectivity, but not in the superhubs ^27^.

**Figure 4:**
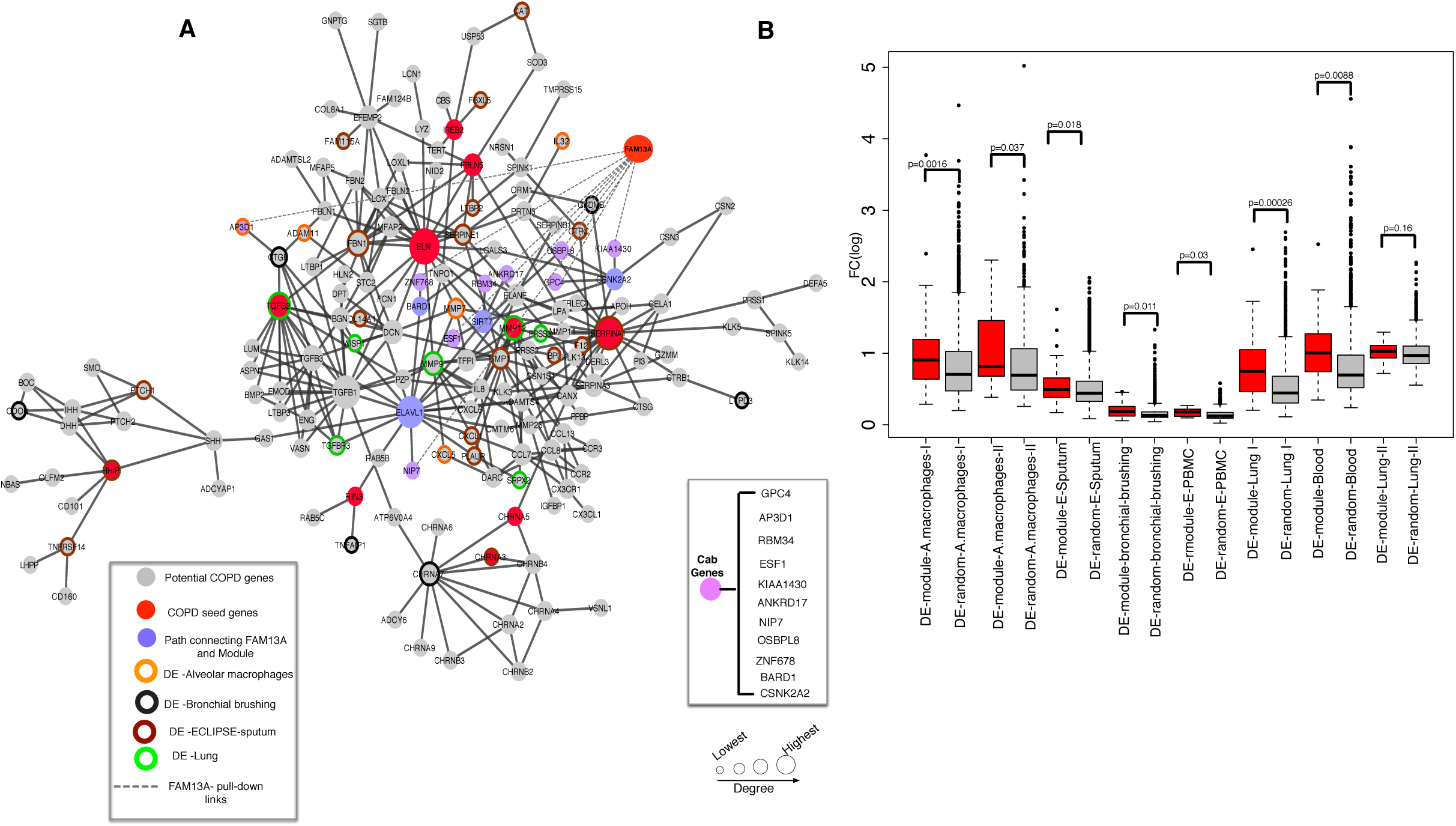
COPD disease network module, including experimentally determined FAM13A interactors, and gene-expression changes in COPD-specific data. **A**. COPD disease network module connecting 11 seed genes including *FAM13A*. **B**. Fold change difference between module differentially expressed genes (p<0.05) and non-module differentially expressed genes.

### COPD disease module with all eleven COPD seed genes

C_AB_ considers all of the possible paths between ⟨CA⟩and ⟨CB⟩genes to calculate the statistical significance; hence, we applied a greedy strategy (Steiner) to find the optimal paths among all of the paths connecting the COPD network neighborhood and C_AB_ genes ^28^. We observed a single network module consisting of C_AB_ genes and COPD network neighborhood genes with only four intermediate genes (*ELAVL1, CSNK2A2, BARD1* and *SIRT7*). Of interest, including these linker genes provided connections to the network module for the two COPD seed genes, *RIN3* and *HHIP*, that were not part of the original 129 genes largest connected component. Our resulting expanded set of 163 connected genes, including all of the 11 seed genes (**Supplementary table 1**), is referred to as the ‘ *COPD disease network module*’ (**Figure 4A**).

**Table 1:** Differentially expressed COPD disease module genes in four datasets with adjusted p-values <0.05

| Shaykhiev2009-Alveolar macrophages |  | Non-smoker vs COPD |  |
| --- | --- | --- | --- |
| Gene.symbol | logFC | adj.P.Val | P.Value |
| IL32 | -3.770 | 0.0014 | 1.35E-06 |
| ADAM11 | -1.441 | 0.0342 | 0.0004 |
| CXCL5 | -2.049 | 0.0359 | 0.0004 |
| MMP7 | 1.952 | 0.0381 | 0.0005 |
| AP3D1 | 0.451 | 0.0387 | 0.0005 |
| MMP12 | 2.390 | 0.0423 | 0.0006 |
| Tedrow2013-Lung |  | Control vs COPD |  |
| Gene.symbol | logFC | adj.P.Val | P.Value |
| MMP1 | 2.452 | 0.0069 | 3.13E-05 |
| TGFB2 | -0.768 | 0.0141 | 0.0001 |
| WISP1 | 1.410 | 0.0158 | 0.0002 |
| PRSS3 | 1.023 | 0.0165 | 0.0002 |
| MMP9 | 1.238 | 0.0240 | 0.0004 |
| TGFBR3 | -0.629 | 0.0298 | 0.0006 |
| CAT | -0.466 | 0.0306 | 0.0007 |
| SRPX2 | 0.728 | 0.0401 | 0.0011 |
| MMP12 | 1.621 | 0.0433 | 0.0013 |
| Singh2011-Eclipse<br>sputum |  | GOLD I vs GOLD IV |  |
| Gene.symbol | logFC | adj.P.Val | P.Value |
| FAM115A | -1.228 | 0.0023 | 2.78E-06 |
| HHIP | -0.645 | 0.0046 | 2.91E-05 |
| CAT | -0.535 | 0.0047 | 3.17E-05 |
| SERPINE1 | 0.969 | 0.0088 | 0.0002 |
| CAT | -0.683 | 0.0100 | 0.0002 |
| CHRNA3 | -0.875 | 0.0109 | 0.0003 |
| MMP1 | 1.320 | 0.0157 | 0.0006 |
| CXCL1 | 0.494 | 0.0167 | 0.0007 |
| TNFRSF14 | 0.492 | 0.0176 | 0.0008 |
| F12 | 0.485 | 0.0184 | 0.0008 |
| LTBP2 | 0.722 | 0.0205 | 0.0010 |
| BPI | 0.870 | 0.0229 | 0.0013 |
| CTRC | 0.625 | 0.0273 | 0.0019 |
| FBN1 | -1.399 | 0.0278 | 0.0019 |

| COL14A1 | -0.410 | 0.0288 | 0.0021 |
| --- | --- | --- | --- |
| SERPINA1 | 0.592 | 0.0294 | 0.0021 |
| FBXL5 | 0.400 | 0.0384 | 0.0035 |
| PLAUR | 0.283 | 0.0394 | 0.0037 |
| PTCH1 | -0.753 | 0.0394 | 0.0037 |
| **Steiling2013-bronchial brushing** |  | **Current smokers NO-COPD-Current smokers with COPD** |  |
| Gene.symbol | logFC | adj.P.Val | P.Value |
| CDON | -0.264 | 0.0296 | 0.0005 |
| LYPD3 | 0.218 | 0.0370 | 0.0009 |
| GSDMB | 0.322 | 0.0431 | 0.0012 |
| CHRNA7 | 0.185 | 0.0438 | 0.0013 |
| CTGF | 0.237 | 0.0477 | 0.0016 |
| TNFAIP1 | 0.126 | 0.0495 | 0.0018 |

### Validation of COPD disease network module in COPD specific gene-expression data

We tested the relevance of the COPD disease network module by evaluating fold change of differentially expressed module genes in COPD-specific gene expression data sets. We compared the fold change (absolute value of logarithm of fold change) of differentially expressed module genes to all other differentially expressed genes with unadj.p<0.05 in eight COPD-specific gene expression data sets (**Table 2**). We observed a significantly higher fold-change in the COPD disease network module compared to other differentially expressed genes in seven datasets (**Figure 4B).** As shown in **Table 2**, even after removing the seed genes, the significance was retained in six datasets (**Supplementary figure 5**). Further, by considering all of the genes tested for differential expression, we still find that COPD disease network module genes were significantly enriched in four COPD gene-expression datasets (sputum, lung tissue, peripheral blood and alveolar macrophages) (**Supplementary figure 6**). These results suggest the ability of our network-based approach to identify new genes relevant to COPD. Additionally, to correct for connectivity as a potential selection bias in the comparison of module and non-module genes, we selected 10 random genes either from the disease network module or from all differentially expressed genes (filtered at p<0.05). For the latter, we made sure that all selected genes were connected using an iterative procedure: the first gene was selected at random, the second gene was selected in the neighborhood of the first gene, the third gene was selected in the neighborhood of the two first genes and so on. As compared to our previous observation in **Supplementary figure 5**, we observed that the selection of a connected subset increases the significance of the differences in gene expression between the COPD disease module genes and randomly selected genes (*p<0.05, **p<0.01, ***p<0.001, **Supplementary figure 7**). This seems to be due to the fact that high fold change genes selected at random when looking at all differentially expressed genes tend to not be connected to other differentially expressed genes. Overall, these results indicate that the differentially expressed genes were heavily localized in the gene set added by our approach, and not influenced by the p-value criteria, thus supporting our method’s ability to identify candidate genes relevant to COPD.

**Table 2:** Enrichment of COPD disease network module genes in different tissue gene expression data sets with and without seed genes

| **Reference** | **GEO ID** | **Tissue** | **P-value with Seed genes** |
| --- | --- | --- | --- |
| Shaykhiev2009 | GSE13896 | Alveolar Macrophages I | 0.002 |
| Poliska2011 | **GSE16972** | **Alveolar Macrophages II** | **0.037** |
| Singh2011 | GSE22148 | Sputum | 0.018 |
| Steiling2013 | GSE37147 | Bronchial brushings | 0.011 |
| Bahr2013 | GSE42057 | Peripheral blood mononuclear cell | 0.030 |
| Tedrow2013 | GSE47460 | Lung homogenate (Lung I) | 0.00026 |
| Singh2014 | GSE54837 | Blood | 0.009 |
| Bhattacharya2009 | GSE8581 | Lung tissue (Lung II) | 0.163 |

**Figure 5:**
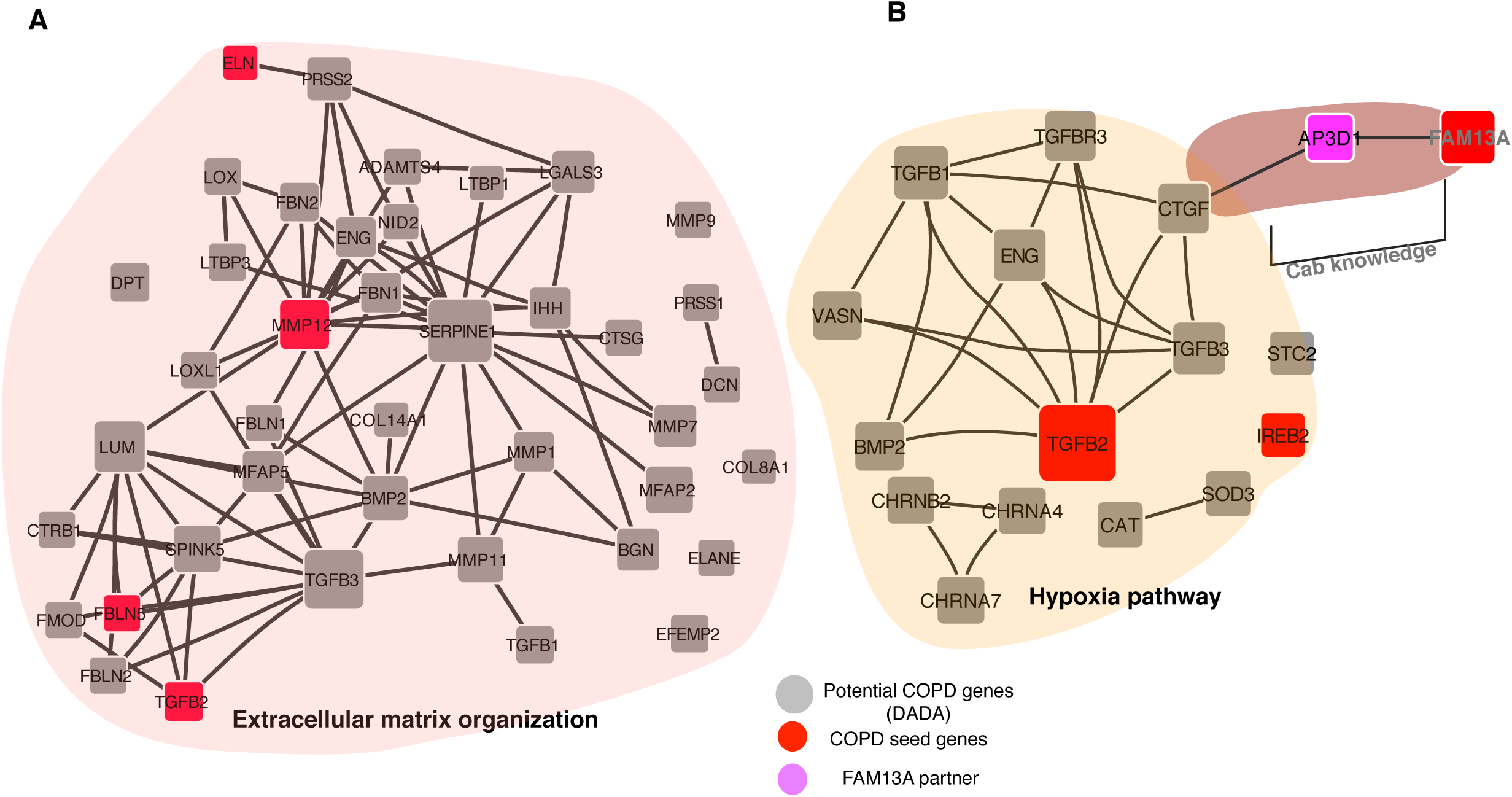
**A.** Extracellular matrix organization pathway genes in COPD disease network module. **B.** Connection of COPD disease network module genes in the hypoxia pathway: ⟨C_AB_⟩helps to connect FAM13A to the hypoxia pathway through CTGF gene.

### Potential candidate genes for COPD

With an adjusted p-value <0.05 (limma), we found 36 COPD disease module genes differentially expressed in different COPD-related datasets. For example, *AP3D1* (adj.p-0.038) and *IL32* (adj.p-0.001) were up-regulated and *MMP12* (adj.p=0.042) was down-regulated in non-smoking controls vs. COPD subjects in alveolar macrophages^29^ (Alveolar macrophage I). In lung tissue, we found *TGFB2* (adj.p=0.014) and *CAT* (adj.p=0.03) were down-regulated in control vs. COPD subjects^30^ (Lung I). Twenty COPD disease module genes were differentially expressed in GOLD stage II vs. GOLD stage IV subjects in ECLIPSE induced sputum data^31^. *CTGF* (adj.p=0.047), *GSDMB* (adj.p=0.044) and *CHRNA7* (adj.p=0.043) were up-regulated between current smokers with no COPD vs. current smokers with COPD in bronchial brushing samples ^32^ (**Table 1**). These results support the ability of our approach to localize candidate genes of potential relevance in COPD-related tissue types. Moreover, all of the 9 C_AB_ genes were differentially expressed in at least one of the gene expression datasets (Z=2.2, p=0.016) (**Supplementary figure 8**).

### Biological pathway enrichment in the COPD disease module

Among the biological pathways most significantly enriched in the COPD disease network module were inflammatory response, collagen catabolic process, regulation of TGFB-receptor signaling pathway, and extracellular matrix organization pathway (**Table 3**). Alterations of extracellular matrix components (ECM), including elastin, are known in patients with COPD, and they contribute to airflow obstruction^33^. In the COPD network module, 34 genes representing the ECM pathway were connected to each other (**Figure 5A**). Moreover, we found support from the medical literature for 23 module genes from the total of 41 genes representing the ECM pathway in COPD pathogenesis (Supplementary table 2). C_AB_ genes were part of: Glycosaminoglycan/aminoglycan catabolic process (*GPC4*), negative regulation of muscle cell differentiation (*ANKRD17*), negative regulation of cell migration (*OSBPL8*), regulation of alpha-beta T cell activation (*AP3D1*) and response to decrease in oxygen levels (*AP3D1*). Gene expression analyses in cell lines from several tissues have demonstrated an increase in FAM13A levels in response to decrease in oxygen levels ^34^. It has been suggested that lower oxygen tension might modulate FAM13A activity^35^, however, the exact mechanism has not been explained. In the COPD disease network module, AP3D1 (C_AB_ gene) interacts with FAM13A and is an immediate neighbor of the CTGF gene, which is part of the hypoxia pathway (decrease in oxygen levels). Thus, the connection of FAM13A to CTGF reveals a potential mechanism by which FAM13A could contribute to the hypoxia response (**Figure 5B)**.

**Table 3:**
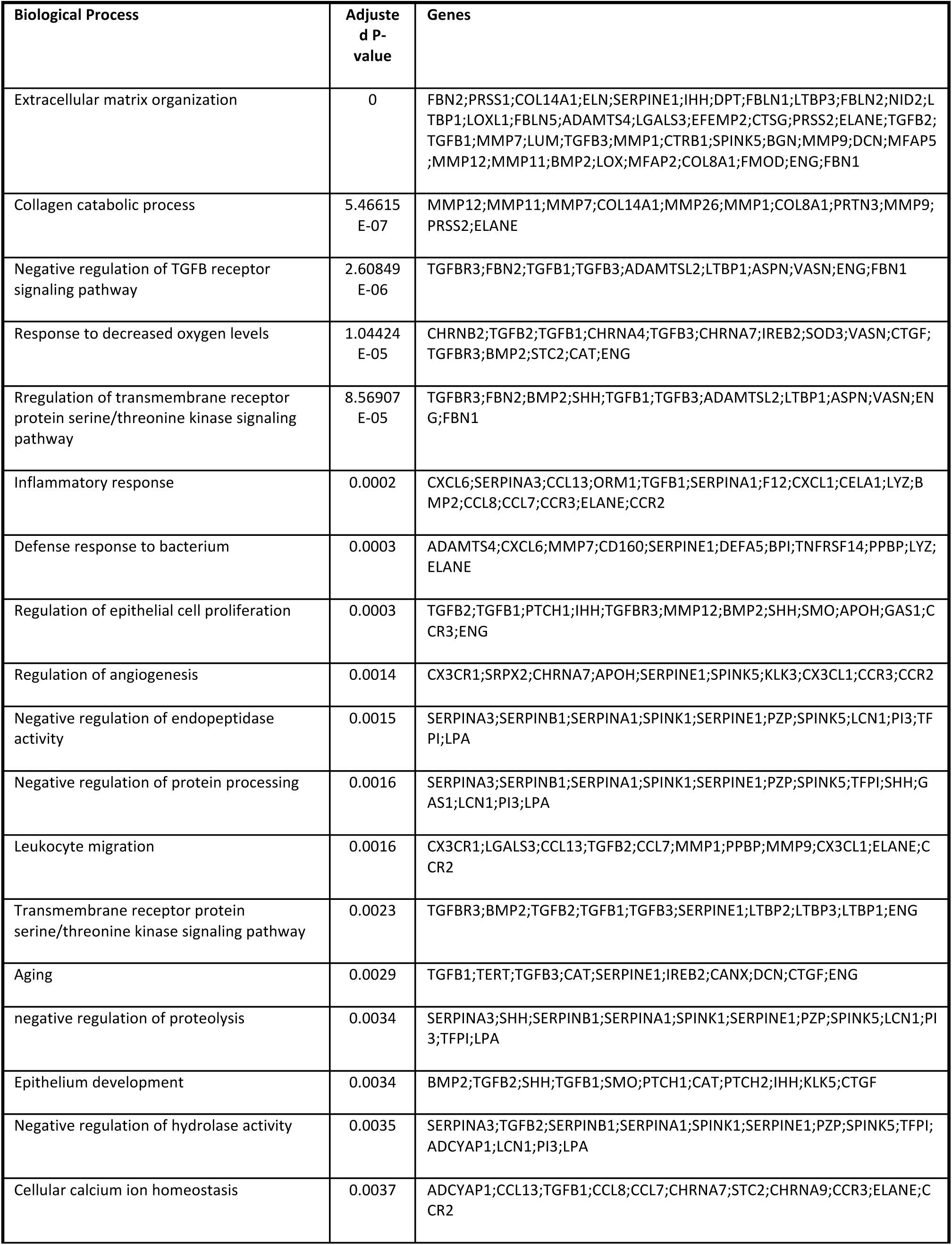
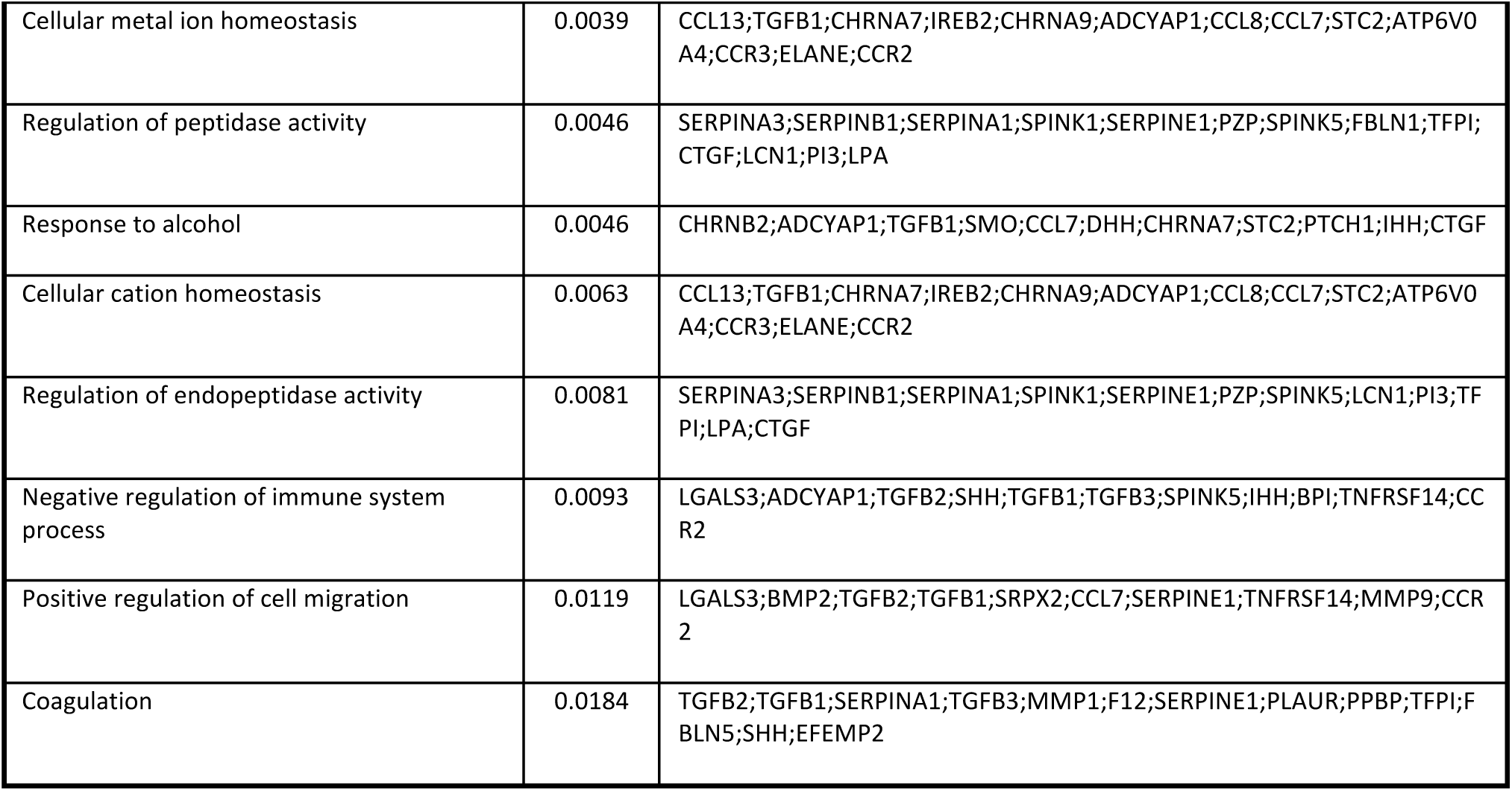
Biological pathways significantly enriched in the COPD disease network module

We observed a small overlap (37 genes, 23%; vs 14% background, p-value=0.0013) of the COPD disease network module with the Inflammasome (see methods) ^36^ (**Supplementary table 1**). This suggests that the COPD disease network module was enriched for inflammation-related genes, which is consistent with the known role of inflammation in COPD ^37^. Overall, the COPD disease network module not only contains the inflammation component, but also other functional components like extracellular matrix organization, hypoxia response, and WNT/beta catenin signaling pathways ^23^.

## Discussion

The purpose of this work was to determine whether a network-based approach could enhance our understanding of the genes involved in the pathogenesis of a complex disease (COPD) by combining new experimental protein-protein interaction data with the existing human interactome. Identifying causal genes for complex diseases like COPD, which are likely influenced by many genetic factors of modest effect size, is a major bottleneck in understanding the biological mechanisms leading to these diseases. A complete and accurate map of the human interactome could have tremendous impact on our ability to understand the molecular underpinnings of human disease. Yet, such a map is far from completion, which makes it currently impossible to evaluate precisely how far a given disease network module is from completion. Here, we showed that despite its incompleteness, a systematic network-based approach could help us to understand the connectivity of disease genes in COPD. Our initial analysis provided a set of 140 potential candidate genes that were part of three connected components in a disease network neighborhood. Interestingly, the largest connected component of this set of genes included 8 of the seed genes, which showed substantial network coherence and localization. Some of these 140 candidate genes have been previously implicated in COPD. For example, *OLFM2* was among the genes within the COPD protein-protein interaction network built with a greedy search algorithm ^38^. *SOD3* is known to attenuate emphysema and reduces oxidative fragmentation of ECM in mouse lung ^39^. In addition, *TGFB1* and its pathway members have been frequently implicated in COPD pathogenesis ^40^. The novel C_AB_ measure assisted in constructing a more comprehensive COPD disease network module including *FAM13A* and its relevant partners, eventually connecting all of the 11 COPD seed genes into a single connected component comprising 163 genes/proteins (**Figure 4A**). Overall, the COPD module genes showed significant differences in gene expression levels from lung tissue, alveolar macrophage, blood, and sputum samples. For example, Tumor necrosis factor alpha (TNFα)-induced protein 1 (*TNFAIP1*) was upregulated in smokers with COPD and directly interacts with RIN3, a COPD GWAS gene ^41^. TNFAIP1 has been reported to be crucial for the induction of apoptosis ^42^, indicating a potential role of RIN3 in apoptosis. Furthermore, *AP3D1* was upregulated in COPD subjects^29^ (Alveolar macrophage I) and directly interacts with FAM13A in our pull-down assay. The new alliance of FAM13A to the COPD disease network module via AP3D1 connects it to the hypoxia pathway (**Figure 5B**), which reveals the potential molecular mechanism by which FAM13A influence hypoxia in epithelial and endothelial cells. Although other lung (e.g., idiopathic pulmonary fibrosis) and heart (e.g., congestive heart failure) diseases can cause hypoxia, hypoxia is a common and important complication of advanced COPD. Thus, we found it interesting that pathway analysis of the COPD network module identified the hypoxia pathway. This evidence suggests the potential of the C_AB_ measure to reveal new disease biology that might have been missed due to the incomplete human interactome.

Our approach adds a new dimension to the current causal gene identification approaches in complex diseases using the human interactome. Moreover, we were able to localize the network neighborhood of COPD and try to address (at least in part) the shortcomings of interactome incompleteness by providing new experimentally derived interactions for *FAM13A*, a key COPD gene not present in the current human interactome. We were able to connect FAM13A individual interactions to a localized network neighborhood by developing a new metric of network closeness, C_AB_. With the current thrust to understand GWAS genes with the help of incomplete protein interaction networks, our approach provides an alternative to connect targeted interaction and interactome data to identify a disease network module.

We focused on only a small set of seed genes for COPD, and that could be one of the limitations of the work. Moreover, since the disease-related gene within each COPD GWAS locus has not been definitively proven, we selected those genes that had the most compelling evidence for a role in COPD pathogenesis. For example, murine models of emphysema have demonstrated a smoking-related phenotypic effect for genes in four of the COPD GWAS loci that we included: 1) *HHIP* ^43^; 2) *FAM13A* ^44^; 3) *IREB2* ^45^; and 4) *MMP12* ^46^. In addition, several other COPD GWAS loci have strong candidate genes, such as the nicotinic acetylcholine receptor genes that have been related to nicotine addiction (*CHRNA3* and *CHRNA5*) and *TGFB2* (part of the TGFBeta pathway). Thus, we contend that most of our selected seed genes are likely related to COPD pathogenesis. We also acknowledge that protein-protein interactions observed during in vitro experiments like yeast two-hybrid or affinity purification assays may not actually occur due to the absence of cellular co-localization or gene expression in the tissue of interest. COPD is a heterogeneous disease, and it is possible that different subtypes of COPD patients could have different disease network modules. Since linker genes connected the three COPD disease components in the COPD network neighborhood into a single disease network module, it could be possible that these are really three different COPD network modules. Thus, future research to identify network modules related to specific COPD subtypes is warranted.

Overall, the disease network module approach that we applied is generic and can be applied to other diseases; this type of approach may be of broad use in disease gene identification in complex diseases in the coming era of network medicine.

## Materials and Methods

### Selection of high confidence COPD-associated genes

Starting with previous GWAS for COPD susceptibility, and with specific genes implicated by eQTL or functional studies within GWAS regions, we identified a set of well-established genes associated with COPD: *HHIP, CHRNA3/CHRNA5/IREB2*, and *FAM13A*. We added recently described genome-wide significant associations to moderate-to-severe COPD or severe COPD, including *RIN3, MMP12*, and *TGFB2* ^41,47-51^. We also considered genes causing Mendelian syndromes which include emphysema as part of their syndrome constellation: alpha-1 antitrypsin deficiency (*SERPINA1*) and cutis laxa (*ELN* and *FBLN5*) ^52,53^. These 11 genes, *in toto*, were subsequently used as seed genes for network analyses. We included several genes from the chromosome 15q25 locus, since previous work from our group has suggested that there are likely at least two COPD genetic determinants in this region—both related to nicotine addiction (nicotinic acetylcholine receptor genes CHRNA3 and CHRNA5) and unrelated to nicotine addiction (IREB2)^54^. Of note, HHIP, FAM13A, and IREB2 are also supported by animal models of emphysema. In addition to these five COPD GWAS genes, we added MMP12, which was associated with COPD before it was discovered by GWAS^55^ and which is also supported by an animal model of emphysema, as well as TGFB2 and RIN3. TGFB2 and RIN3 (as well as HHIP, FAM13A, and the chromosome 15q25 region) were also strongly supported by the recent International COPD Genetics Consortium GWAS^7^.

### Human protein interaction network: Interactome

We compiled the physical protein-protein interactions from the ConsensusPathDB database ^56^. Physical protein interactions were assigned a confidence score between 0 and 1 using the interaction confidence-scoring tool (IntScore)^57^. We relied only on physical interaction data in ConsensusPathDB, obtaining *M=*150,168 links between *N=*14,280 genes encoding these proteins with mean degree <k> of 21.03 and average clustering coefficient <C> of 0.141.

### Localization of COPD network neighborhood in the human interactome

The concept that proteins located close to one another in the human interactome may cause similar diseases is becoming an increasingly important factor in the search for complex disease genes. Different approaches tackle this problem of predicting complex disease susceptibility genes using different kinds of integrative data, but all of them involve superimposing a set of candidate genes alongside a set of known disease genes in some physical or functional network ^13,17,58,59^. However, many existing methods are likely to favor highly connected genes, making prioritization sensitive to the skewed degree distribution of protein-protein interaction (PPI) networks, as well as ascertainment bias in available interaction and disease association data. To enhance our understanding regarding the local neighborhood of seed genes in the network, we applied the degree aware algorithm (DADA) ^20^ to compute the proximity of the selected COPD seed genes to their neighbors by exploiting the global structure of the network. Several studies ^17,60^ have shown global approaches like random walk outperform other local approaches like shortest path distances, and therefore we focused on the global method. The final ranking for 14,280 genes encoding proteins included in the network was achieved by merging the random walk restarts output and statistical adjustment models. We used the results from a COPD GWAS of 6,633 cases and 5,704 controls from 4 cohorts to define a boundary for the most promising DADA-ranked genes ^41^. We assigned significant SNPs to genes using 50kb boundaries, and generated gene-based p-values using VEGAS ^21^.

### FAM13A pull-down assay

The *FAM13A* gene resides at a locus associated with COPD and with lung function in the general population by GWAS ^41,51,61,62^. *FAM13A* contains a Rho GTPase-activating protein-binding domain, inhibits signal transduction, and responds to hypoxia; however, its primary function in the lung remains to be determined. The pull-down assay using affinity purification-mass spectrometry was performed previously^23^ and resulted in 96 interacting proteins, establishing 96 edges for *FAM13A* in the interactome.

### Proximity of the targeted interactions to the COPD neighborhood-Cab

To quantify the network-based separation between the identified *FAM13A* interactions and the COPD disease network neighborhood, we introduce the Cab minimum weighted distance, which we define as follows:

For any two nodes *l* and *m* we define the Cab distance as:

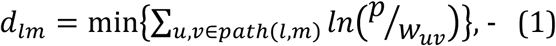

where *w*_*uv*_ ∈ [0, 1] is the edge confidence score and p ∈ (1, ∞) is the parameter of the model. Note that distance *d*_*lm*_ depends both on the total number of network-based edges one needs to traverse from node *l* to node *m* and also the confidence scores of these weights, while parameter *p* tunes the relative contribution of these two factors.

In particular, in the *p* = 1 case, *d*_*lm*_ depends only on confidence scores of edges connecting two nodes:

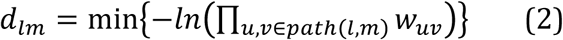

If confidence scores *w*_*uv*_ are regarded as independent probabilities for the edges to be present in the network, then the product in Eq. (2) is simply the probability that given path from *l* to m exists. The larger this probability, the smaller distance *d*_*lm*_ is. On the other hand, if *p* is large, then *d*_*lm*_ is independent of confidence scores:

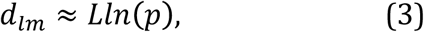

where *L* is the smallest number of edges that need to be traversed from *l* to m. We then use *d*_*lm*_ to define distance from *l* to a set of nodes *M* as the sum of distances from *l* to all nodes in *M*:

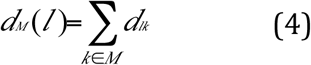

We evaluated distances from nodes to the neighborhood for a set of parameters *p* = {*e*^0^, *e*^1^, *e*^2^, *e*^10^}. In the following, the values of the parameter are indexed with power of the exponent (0,1,2,10). To quantify the significance of the observed distribution of distances *P =* (*d*_*lm*_) from target proteins to the COPD localized neighborhood we used the Mann–Whitney U test with significance cutoff of P < 0.05. Specifically, we calculated the distribution of distances between targeted proteins to the module *P* = (*d*_*lm*_) and a random distribution of distance from target proteins to all proteins in the network *P =* (*d*_*lm*_). To measure how much the two distributions are different, we calculate the Z-score:

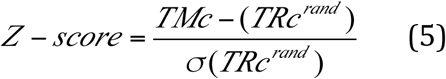

Where *TRc*^*rand*^and *σ*(*TRc*^*rand*^) denote the mean value and standard deviation of the random expectation *p*^*rand*^(*TRc*). Assuming normality of *p*^*rand*^(*TRc*), we can analytically calculate a corresponding p-value for each z-score, yielding a threshold of z-score ⩽ −1.6 for the distance to be smaller than expected by chance with significant p-value ≤ 0.05.

### Local Radiality (LR) method for target prediction

The LR method quantifies the proximity of a node from a set of genes of interest. The LR score of node n in the network G is calculated as follows ^26^:

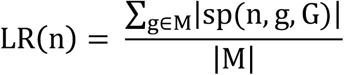

Where sp calculates the length of shortest path between nodes n and g. and |M| is the size of the community of interest (150 genes).

In other words, LR calculates the average shortest paths from node n to the module M.

### COPD network module overlap with inflammasome genes

Since clinical COPD is influenced by inflammation,^63^ we looked for the potential overlap between the COPD disease network module and recognized genes relevant to inflammatory response or the ‘inflammasome’ genes. These inflammasome signature genes were compiled from 11 disease models (asthma, COPD, fibrosis, atherosclerosis, diabetes (adipose), diabetes (islet), obesity, stroke, neuropathic pain, inflammation pain and sarcopenia) ^36^. We used the total of 2,483 inflammatory signature genes previously reported from mouse models and converted them to their human orthologs, obtaining 2,331 genes in our analysis. Mouse to human orthologs were extracted from the Mouse Genome Informatics (MGI) database (http://www.informatics.jax.org).

### Validation of COPD disease module in COPD-specific gene-expression data

Our disease network module approach selects genes based on their topological closeness to the COPD seed genes. To evaluate COPD-specific relevance of genes localized around the seed genes, we extracted significantly differentially expressed genes (p-value<0.05) from eight publicly available COPD-specific gene-expression datasets and assessed for each case the fold change difference between genes present in the COPD disease module compared to non-module differentially expressed genes. We used the limma R package (ver 3.10.1) for differential expression analysis. The 8 datasets are as follow:

1. *Singh2014*: Peripheral blood gene expression samples from 171 subjects from the Evaluation of COPD Longitudinally to Identify Predictive Surrogate Endpoints (ECLIPSE) study (GSE54837). Differential expression analysis was performed between control (n=6) (healthy nonsmokers) vs. severe COPD (n=13)^64^.

2. *Singh2011*: Induced sputum gene expression from 148 COPD subjects in the ECLIPSE study, with 69 Global Initiative for Chronic Obstructive Lung Disease (GOLD) stage 2, and 71 GOLD stage 3 & 4 subjects (GSE22148). Gene expression differences between GOLD 2 and GOLD 3&4 were analyzed ^31^.

3. *Shaykhiev 2009*: Transcriptional profiling of alveolar macrophages obtained by bronchoalveolar lavage of 24 healthy nonsmokers and 12 COPD smokers (GSE13896) ^29^.

4. *Bahr2013*: Expression data from peripheral blood mononuclear cells (PBMC) generated from 136 subjects from the COPDGene study (*GSE42057*), which consisted of 42 ex-smoking control subjects and 94 subjects with varying severity of COPD ^65^.

5. *Tedrow2013*: Microarray data from whole lung homogenates of subjects undergoing thoracic surgery from the Lung Tissue Research Consortium (LTRC). These subjects were diagnosed as being controls or having COPD as determined by clinical history, chest CT scan, and surgical pathology. We considered 220 COPD subjects and 108 controls with no chronic lung disease by CT or pathology. These subjects went for surgery typically to investigate a pulmonary nodule and normal lung tissue was obtained for differential expression analysis (*GSE47460)* ^*30*^.

6. *Bhattacharya2009*: Gene expression patterns in lung tissue samples derived from 56 subjects (*GSE8581)*. Cases (n=15) were defined as subjects with FEV1<70% predicted and FEV1/FVC<0.7 and Controls (n=18) as subjects with FEV1>80% predicted and FEV1/FVC>0.7 ^66^.

7. *Poliska2011:* Gene expression data from alveolar macrophage samples from 26 COPD and 20 healthy control subjects (GSE16972) ^67^.

8. *Steiling2013*: Bronchial brushings obtained from current and former smokers with and without COPD (GSE37147). Data from 238 subjects was used in the analysis to determine the association of gene expression with COPD-related phenotypes ^32^.

## Declarations

### Acknowledgments

We thank Craig Hersh, Dawn Demeo, and Jarrett Morrow for the discussion regarding the gene expression data. We would like to thank Arda Halu for his help and discussion on protein-protein interaction data analysis. The work was supported by the NIH/NHLBI grants: P01 HL105339, R01 HL111759, P01 HL114501, U01 HL089856 (EKS); CEGS-P50 HG004233-06, R01 HL118455 and R37 HL066289; U01 HL089897 (JDC) and R01HL113264 (E.K.S. and M.H.C.)

We thank Amund Gulsvik, Per Bakke, Augusto Litonjua, Pantel Vokonas, Ruth Tal-Singer, and the GenKOLS, NETT/NAS, ECLIPSE, and COPDGene studies for use of GWAS meta-analysis data.

The COPDGene study (NCT00608764) was funded by U01 HL089856 and U01 HL089897 and also supported by the COPD Foundation through contributions made to an Industry Advisory Board comprised of AstraZeneca, Boehringer Ingelheim, GSK, Novartis, Pfizer, Siemens and Sunovion. The National Emphysema Treatment Trial was supported by the NHLBI N01HR76101, N01HR76102, N01HR76103, N01HR76104, N01HR76105, N01HR76106, N01HR76107, N01HR76108, N01HR76109, N01HR76110, N01HR76111, N01HR76112, N01HR76113, N01HR76114, N01HR76115, N01HR76116, N01HR76118 and N01HR76119, the Centers for Medicare and Medicaid Services and the Agency for Healthcare Research and Quality. The Normative Aging Study is supported by the Cooperative Studies Program/ERIC of the US Department of Veterans Affairs and is a component of the Massachusetts Veterans Epidemiology Research and Information Center (MAVERIC). The Norway GenKOLS study (Genetics of Chronic Obstructive Lung Disease, GSK code RES11080), the ECLIPSE study (NCT00292552; GSK code SCO104960), and the ICGN study were funded by GlaxoSmithKline.

## Competing interests

The authors declare the following competing interests. In the past three years, Edwin K. Silverman received honoraria and consulting fees from Merck, grant support and consulting fees from GlaxoSmithKline, and honoraria and travel support from Novartis.

## Author’s contributions

AS and MK conceived the idea for this study and EKS and MHC co-supervised the analyses. AA, JM, XZ, ZJ and MS performed the computational, statistical analyses and experiments. AA, EKS, JC, THB, PSB contributed to the interpretation of the results and in writing of the paper. All authors have read and approved the final manuscript.

---

**Supplementary Figure 1.**
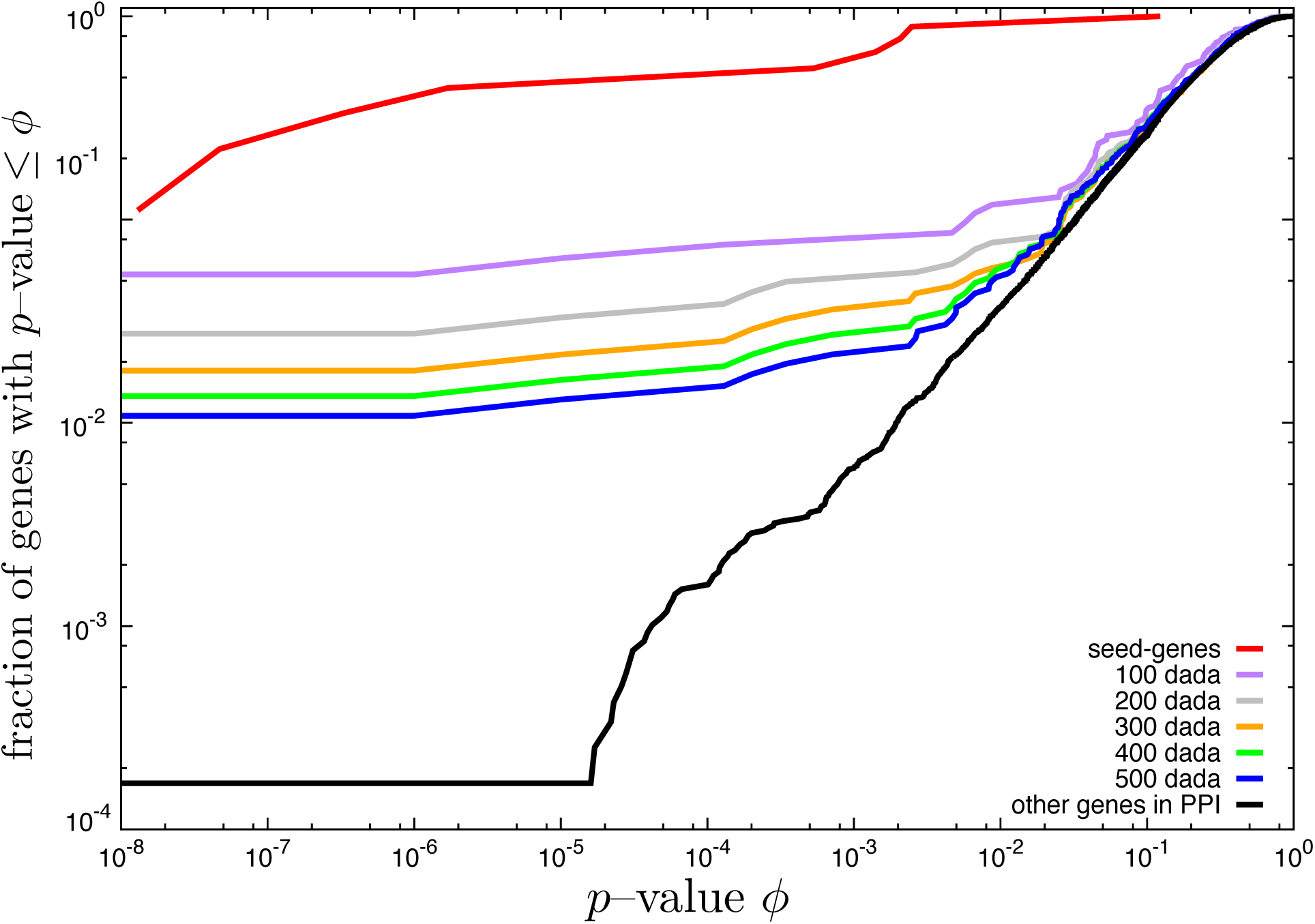

**Supplementary Figure 2.**
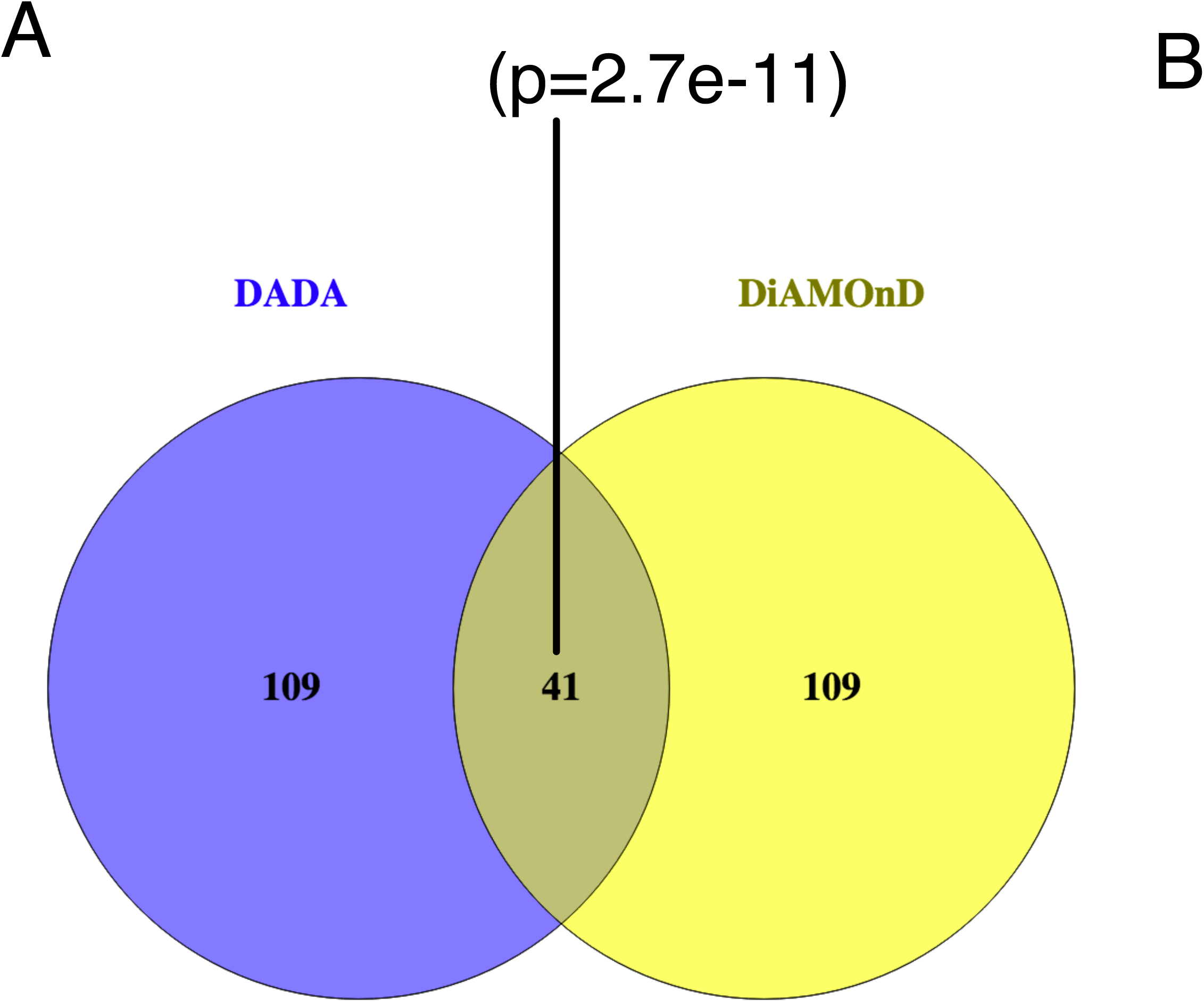

**Supplementary Figure 3.**
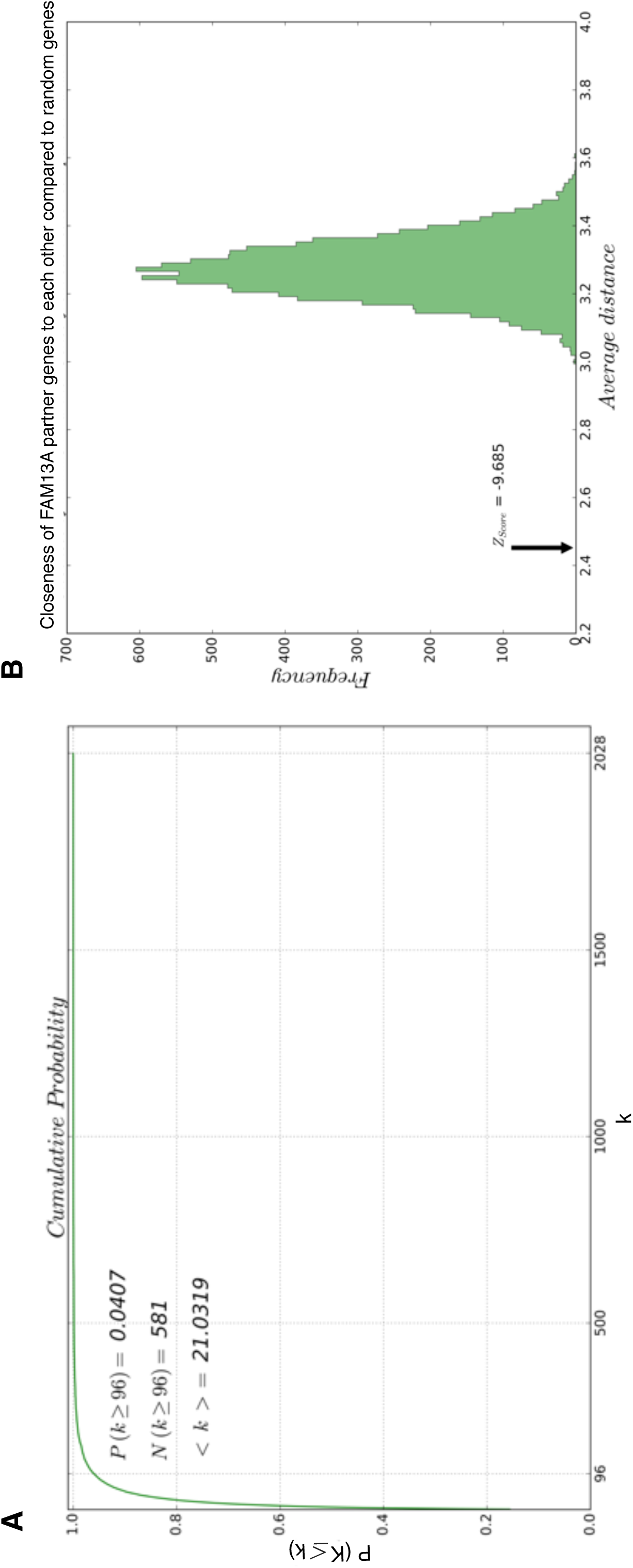

**Supplementary Figure 4.**
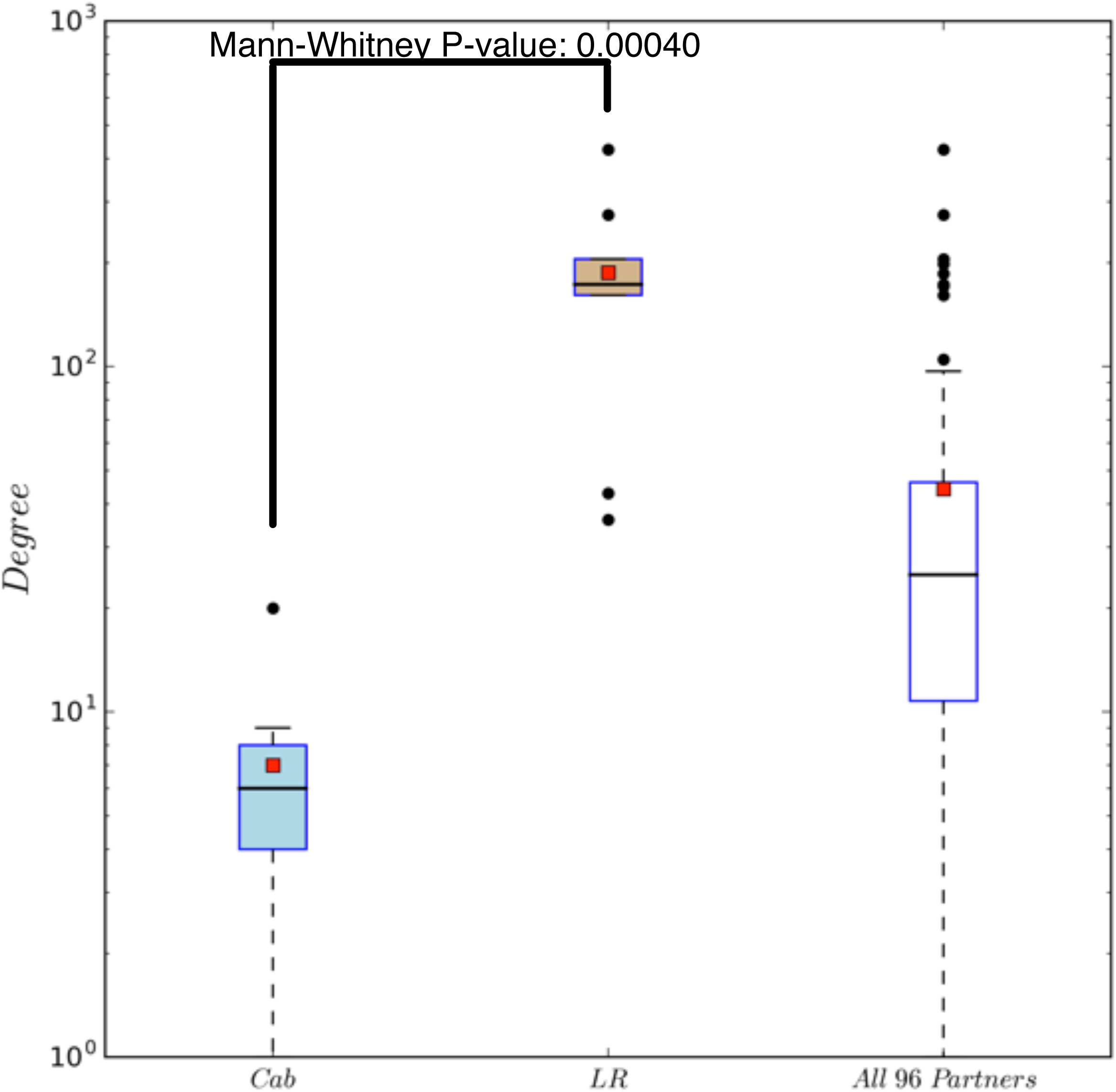

**Supplementary Figure 5.**
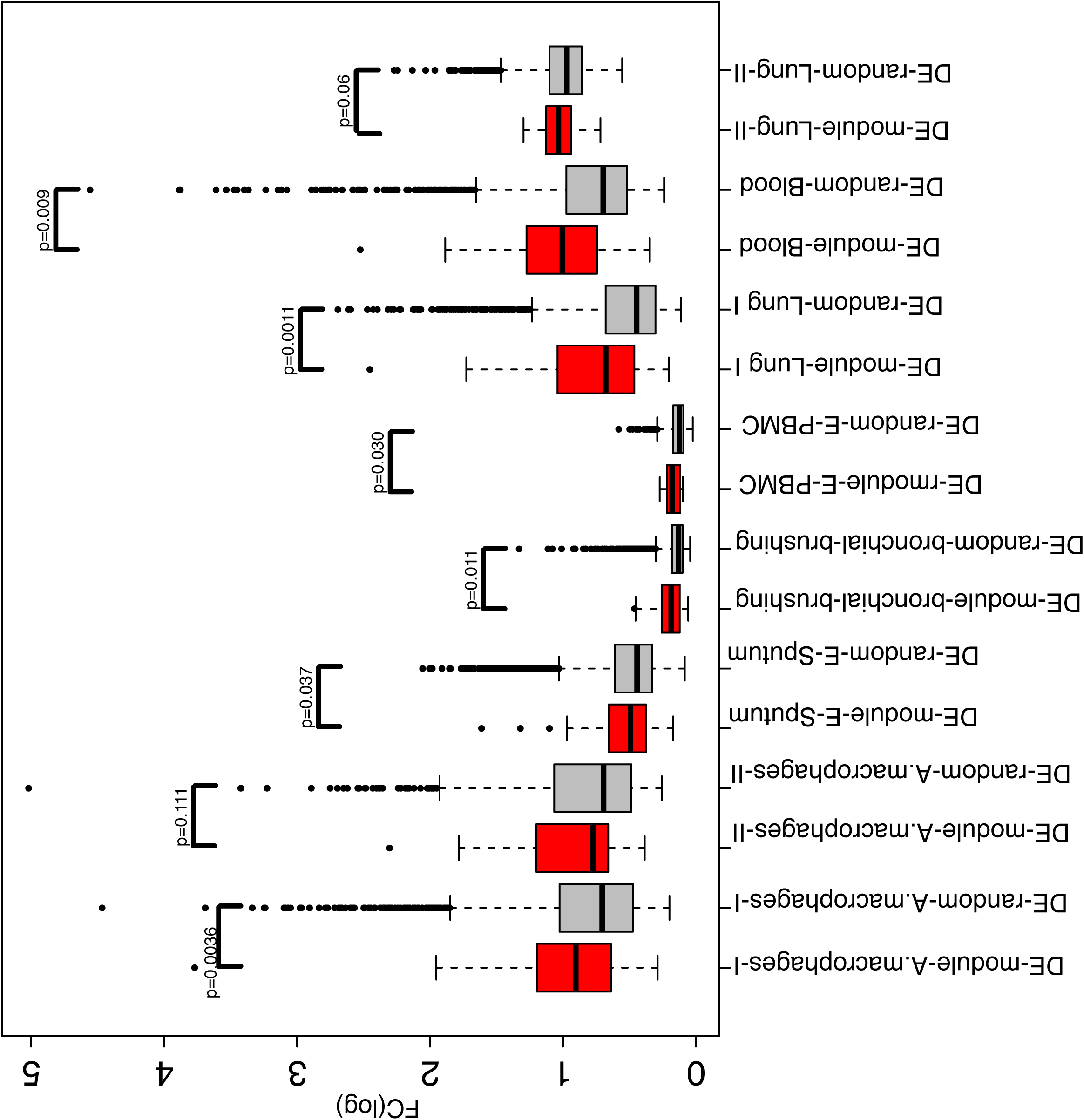

**Supplementary Figure 6.**
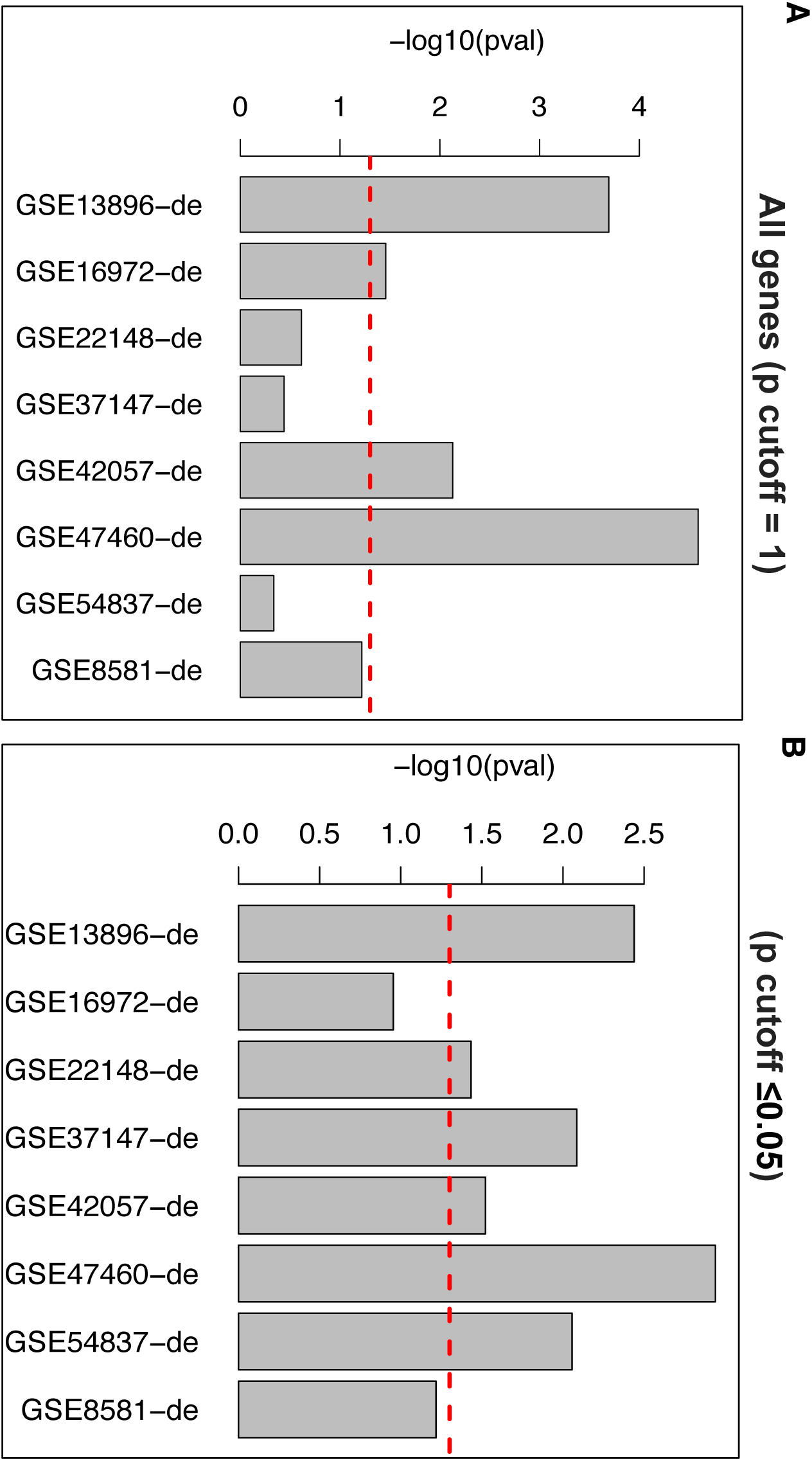

**Supplementary Figure 7.**
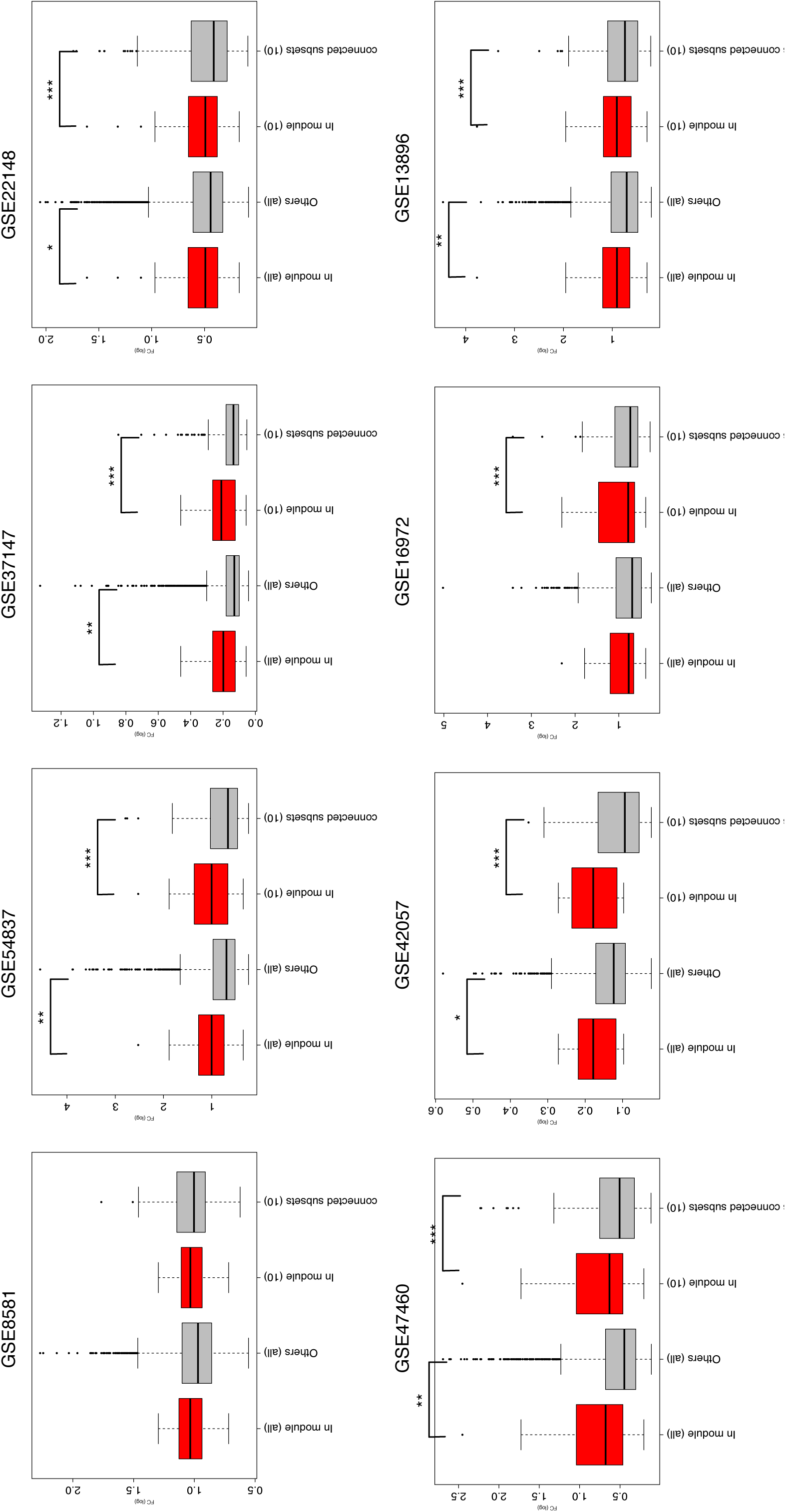

**Supplementary Figure 8.**
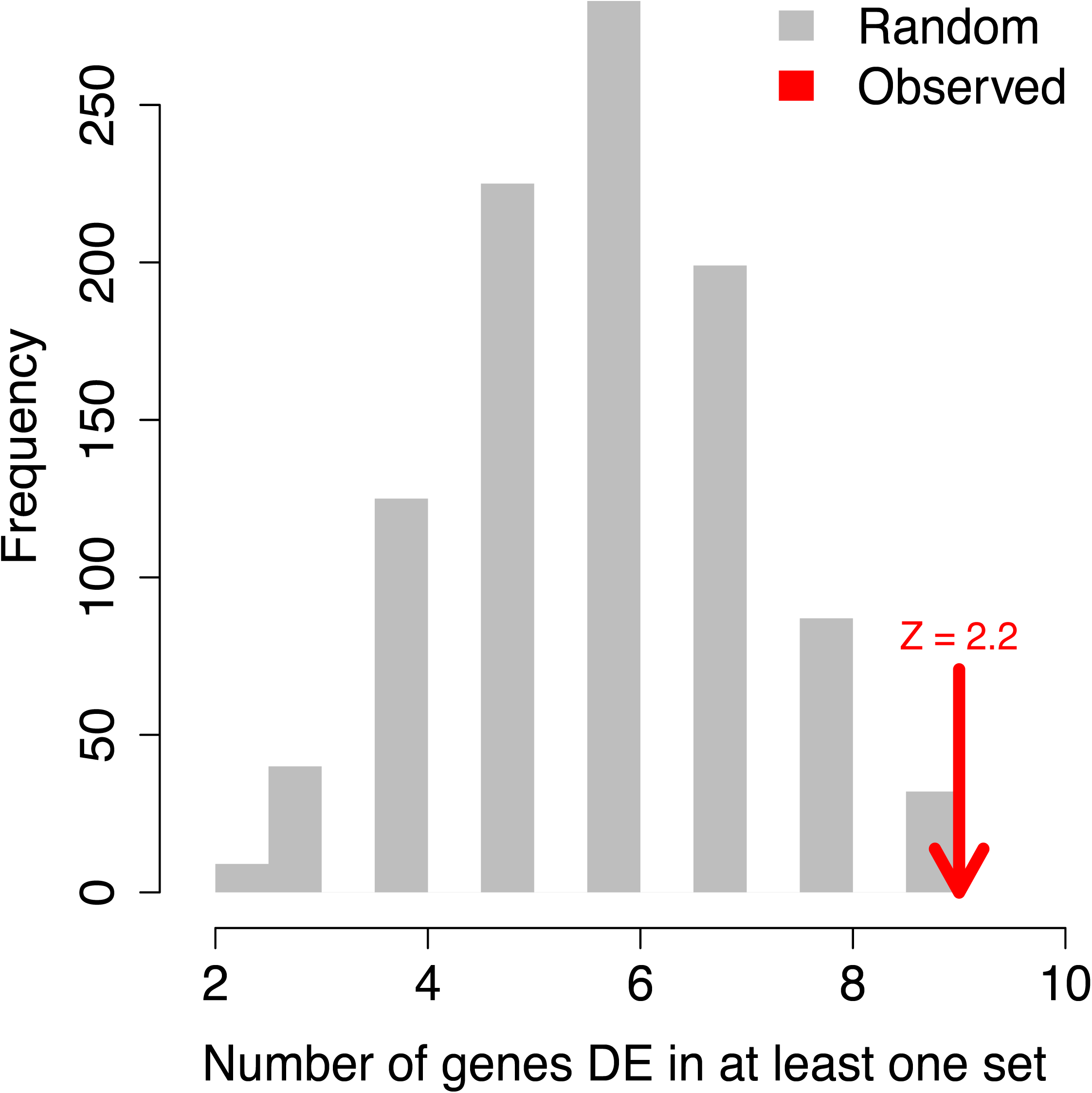

